# Subnanometer structure of medusavirus capsid during maturation using cryo-electron microscopy

**DOI:** 10.1101/2024.03.07.583866

**Authors:** Ryoto Watanabe, Chihong Song, Masaharu Takemura, Kazuyoshi Murata

**Author notes:** Corresponding author and Lead contact: Kazuyoshi Murata. Present address: Core Research Facility, Pusan National University, Yangsan 50612, Republic of Korea (Chihong Song).

## Abstract

Medusavirus is a giant virus classified into an independent family of *Mamonoviridae*. Medusavirus-infected amoebae release immature particles in addition to the virions. These particles were suggested to exhibit the maturation process of this virus, but the structure of these capsids during maturation remains unknown. Here, we applied a block-based reconstruction method in cryo-electron microscopy (cryo-EM) single particle analysis to these viral capsids, extending these resolutions to 7-10 Å. The map revealed a novel network consisted of minor capsid proteins (mCPs), supporting major capsid proteins (MCPs). A predicted molecular model of the MCP fitted into the cryo-EM map clarified the boundaries between the MCP and the underlining mCPs and between the MCP and the outer spikes, identifying the molecular interactions between the MCP and these components. Several structural changes of mCPs were observed beneath the 5-fold vertices of immature particles, depending on the presence or absence of the underlying internal membrane. Furthermore, the lower part of penton proteins on the 5-fold vertices were also lost in the mature virions. These dynamic structural changes of mCPs exhibit important functions in the maturation process of medusavirus.

**Importance:** Structural changes in giant virus capsids during maturation have not been known. Medusavirus is a unique giant virus, where amoebae infected with the virus release immature particles in addition to mature virions. In this study, immature and mature medusavirus particles were investigated using cryo-electron microscopy, and firstly reported the structural changes in the viral capsid during maturation. In DNA empty particles, the conformation of the minor capsid proteins dynamically changed depending on the presence or absence of the underlying internal membranes. In DNA full particles, the lower part of the penton proteins was lost. These are the first report of the structural changes in the viral capsid during giant virus maturation.

## Introduction

Medusavirus is one of the nucleocytoplasmic large DNA viruses (NCLDVs), which was first discovered from a hot spring water in Japan (1). As the characteristic feature, the genome of medusavirus contains a complete set of histone proteins encoding four core histones and one linker histone, as well as a DNA polymerase located at the root of the eukaryotic clade. The facts suggest that medusavirus is phylogenetically more closely related to eukaryotes than other NCLDVs. Medusavirus was, therefore, independently classified as initially *Medusaviridae* and later reclassified as *Mamonoviridae* in the phylum *Nucleocytoviricota* (2). However, a sister strain of medusavirus, *Medusavirus stheno*, was later isolated from a river in Kyoto, Japan (3), indicating that medusavirus is more widely distributed.

Medusavirus also exhibits a unique feature with respect to the replication system and the particle structure. In our previous study, four types of medusavirus particles (pseudo-empty, empty, semi-full and full DNA particles) were identified in the culture medium of amoebae that are infected by medusavirus (4). Time course observations of medusavirus-infected amoeba cells by conventional-transmission electron microscopy demonstrated that these four types of particles exhibit the viral particle maturation process. In the initial stage of infected ameba cell, pseudo-DNA-empty particles appeared in the cytoplasm, and the materials inside the particle are released to form the DNA-empty particles. Then, they are started to be filled with viral DNA (semi-DNA-full particle). Finally, the capsid is completely filled with viral DNA (DNA-full particle), and the matured DNA-full particles are released together with the immature particles (pseudo-DNA-empty, DNA-empty and semi-DNA-full) to the outside of the cell. These medusavirus particles commonly show an icosahedral capsid of T=277 with a diameter of approximately 260 nm, covered with spikes approximately 14 nm long (1, 4). The cryo-EM single-particle analysis (SPA) elucidated their structures at 19.5 Å and 21.5 Å resolutions for the DNA-full and the DNA-empty particles, respectively. The DNA-full particle was approximately 1 nm smaller than the DNA-empty particle, but these particles structurally look similar for the capsids at these resolutions.

Several icosahedral giant viruses have also been studied for their capsid structures using cryo-EM SPA. The icosahedral capsids are commonly composed of a combination of 12 pentasymmetrons and 20 trisymmetrons (5). For the capsid formation of the icosahedral giant viruses, a spiral assembly pathway has been proposed, in which particle assembly initiates in a spiraling fashion around the 5-fold vertices, based on the structural orientation of the MCP capsomers on the pentasymmetron (6). PBCV-1, ASFV, and SGIV have been successfully reported at near-atomic resolution using cryo-EM SPA, identifying not only MCPs but also minor capsid proteins (mCPs) supporting MCPs (7–9). In these image processing, block-based reconstruction method was commonly used (10), where such large objects with a big defocus gradient can be split into several smaller blocks based on the symmetry axes on the icosahedral capsid, and independently reconstructed in three-dimensions (3D). Resultantly, the resolution limit paused by Ewald sphere effect can be extended. The higher resolution map revealed that mCPs form a unique molecular network under the MCP layer, supporting the MCP array directly and the viral DNA via the internal membrane. In PBCV-1, mCPs named P3, P4, and P5 form a trapezoidal unit, which further forms the large trisymmetron. In the giant viruses of *Marseilleviridae*, the trapezoidal units were connected through another mCP named “cement component” to form the trisymmetron (11, 12). In contrast, the trisymmetron of ASFV was constructed with a mCP network consisting of a single mCP (p17) (13).

In this study, we present the capsid structure of medusavirus at 7.3-9.9 Å resolution by using cryo-EM SPA and block-based reconstruction method. The cryo-EM map clarified the boundaries between the MCP and the inner mCPs and between the MCP and the outer spikes. Potentially interacting residues for these connections are identified in the MCP. We also identified a medusavirus-specific mCP network, which is different from the known mCP networks of other giant viruses. Under the 5-fold axis, a conformational change of mCPs in pentasymmetron was further observed with and without internal membrane, where a part of the penton protein was lost in DNA-Full particles, when it compares with that of DNA-Empty particles. These results suggest conformational changes of the capsid proteins during the virion formation, and propose a new model of the maturation process of giant viruses.

## Results

### Subnanometer resolution structures of DNA-Empty and Full particles

In our previous study, the structure of medusavirus particle was reconstructed at 21.5 Å and 19.5 Å resolution by using 4,551 DNA-Empty and 6,981 DNA-Full particles and imposing icosahedral symmetry, respectively (4). In this study, we applied block-based reconstruction method for these cryo-EM data, where the images around the 5-, 3-, 2-fold symmetry axis of the icosahedral particles were independently extracted and reconstructed at 3D. The resolution limitated by defocus gradient were extended to 7.3-9.9 Å (Table 1, Fig. S1, Fig. 1A-C). The entire view of the medusavirus particle was built by merging each block (Fig. 1D). In the 5-fold block, the maps were classified with their structures of the presence or absence of the internal membrane (IM), in addition to the DNA-Full and DNA-Empty particles (Fig. S1C and D).

**Table 1.** Cryo-EM data set.

|  |  |  |
| --- | --- | --- |
| Microscope | Titan Krios G3 |  |
| Accelerating voltage (kV) | 300 |  |
| Spherical aberration (mm) | 0.1 (Cs corrector) |  |
| Detector | Falcon III |  |
| Total dose (e<sup>-</sup>/Å<sup>2</sup>) | 30 |  |
| Micrographs | 2,084 |  |
| Frames per micrograph | 40 |  |
| Nominal magnification | 22,500× |  |
| Pixel spacing on the specimen (Å/pixel) | 3.03 |  |
|  | DNA-Empty | DNA-Full |
| Initial particles picked | 4,625 | 7,038 |
| Final particles used | 4,551 | 6,981 |
| Symmetry imposed | I1 | I1 |
| Resolution (Å) | 21.5 | 19.5 |

**Table 2.** Reconstruction details.

|  | 5-fold block |  |  | 3-fold block |  | 2-fold block |  |
| --- | --- | --- | --- | --- | --- | --- | --- |
| Particle types | DNA-Empty |  | DNA-Full | DNA-Empty | DNA-Full | DNA-Empty | DNA-Full |
|  | with IM | without IM |  |  |  |  |  |
| Particles | 19,028 | 21,900 | 67,456 | 127,482 | 148,795 | 208,834 | 189,286 |
| Symmetry | C5 | C5 | C5 | C1 | C1 | C1 | C1 |
| Resolution | 8.62 | 8.81 | 7.32 | 9.93 | 8.54 | 9.88 | 8.92 |

**Table 3.** Refinement and validation statistics.

|  |  |
| --- | --- |
| Map | 3-fold block (DNA-Full) |
| Initial model | AlphaFol2 generated model |
| R.m.s. deviations |  |
| Bond lengths (Å) | 0.004 |
| Bond angles (°) | 0.983 |
| Validation |  |
| MolProbity score | 2.41 |
| Clashscore | 25.60 |
| Ramachandran plot (%) |  |
| Favored | 91.12 |
| Allowed | 8.57 |
| Outliers | 0.31 |

**Figure 1.**
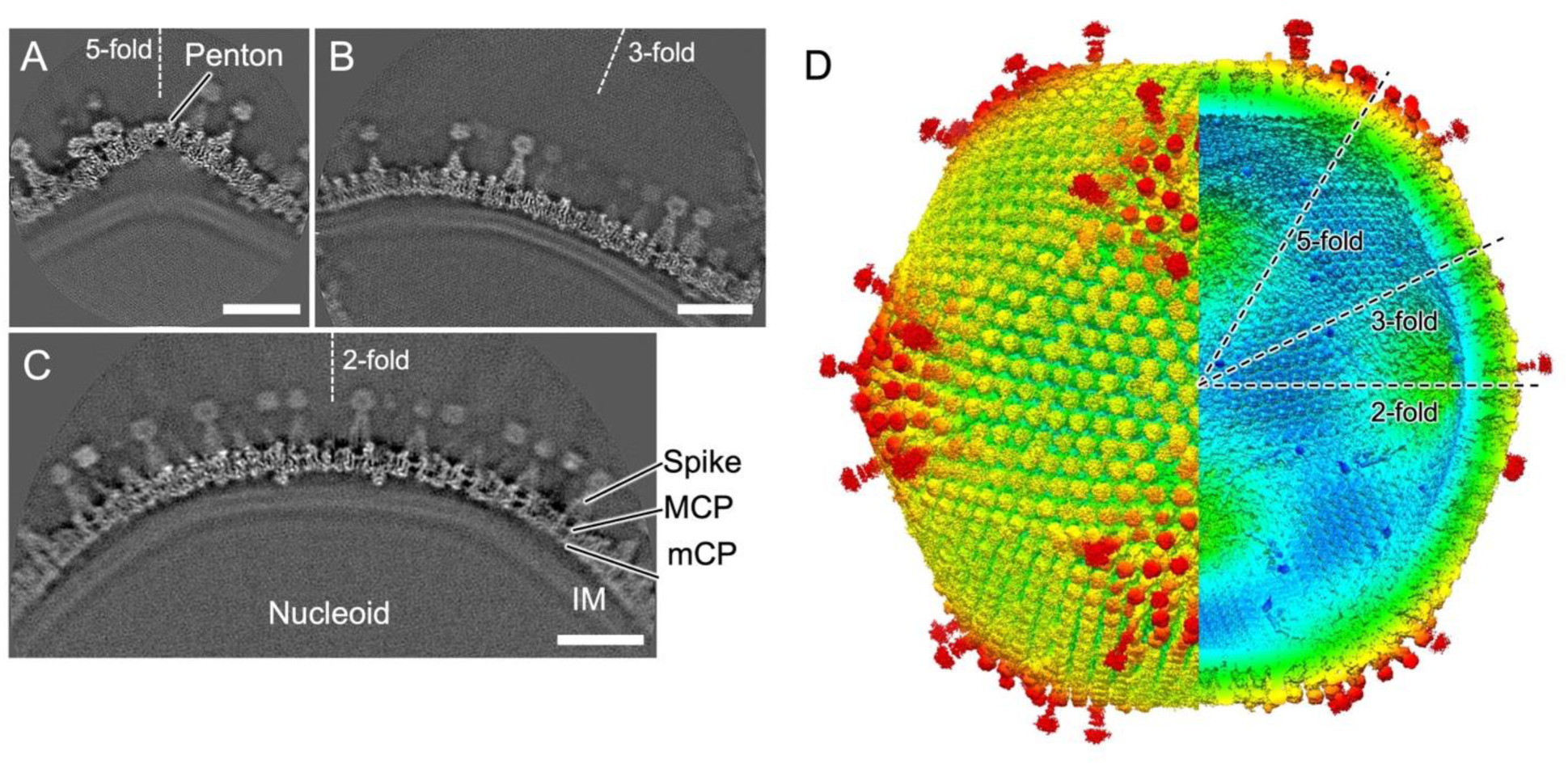
Subnanometer resolution structure of medusavirus capsid. (A, B, and C) Each slice of the block-based reconstruction maps centered on the 5, 3 and 2-fold axes. Under the capsid, the internal membrane (IM) wraps around the nucleoid. The capsid consists of major capsid proteins (MCPs), spikes, minor capsid proteins (mCPs), and pentons on the 5-fold axis. Scale bars = 20 nm. (D) Entire medusavirus capsid merged with each capsid map individually generated by block-based reconstruction. The map is colored by radius. The icosahedral 5, 3 and 2-fold axes are indicated as dotted line.

### The network structure of minor capsid proteins

The mCPs form protein unique networks under the MCP array in the pentasymmetron and trisymmetron, respectively (Fig. 2A). Here, these proteins were named as components based on their structural features, because they have not been identified in each protein sequence at the current resolutions. In the pentasymmetron, the penton proteins located at the center of the 5-fold vertices, were easily identified (purple in Fig. 2A, B). In addition, mCPs were segmented into four components, named PC-I, PC-II, PC-III, PC-IV (Fig. 2B left). PC-I is located near the penton and supports the array of the MCPs labeled P1 (turquoise green in Fig. 2B left). PC-II is located around these and supports the array of other MCPs labeled P3 to P5 (salmon pink in Fig 2B left). PC-III and PC-IV were filamentous components located under these mCP networks and functions to connect trisymmetron to pentasymmetron (light purple and blue in Fig. 2 left).

**Figure 2.**
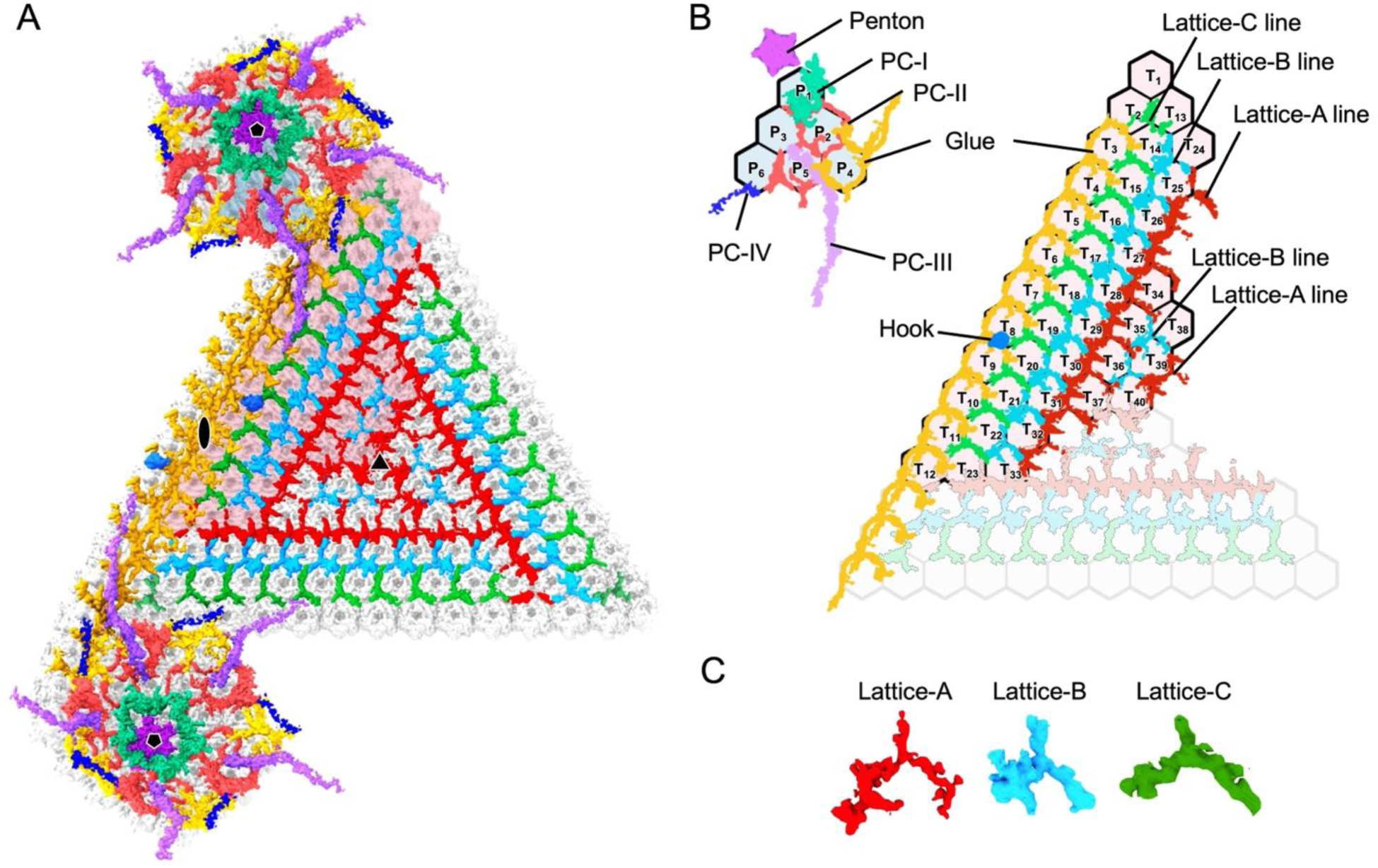
Minor capsid protein network in medusavirus capsid. (A) Cryo-EM map of the minor capsid proteins and the penton, viewed from the interior of the pentasymmetron and trisymmetron. The 5-fold, 3-fold, and 2-fold axes are indicated by pentagon, triangle, and ellipses, respectively. (B) Schematic drawing of the minor capsid proteins and pentons viewed from inside the capsid. Each minor capsid protein component including the penton, is colored according to the shapes as follows: penton (purple), PC-I (turquoise green), PC-II (salmon pink), PC-III (light purple), PC-IV (blue), Lattice-A (red), Lattice-B (light blue), Lattice-C (green), Glue (yellow), and Hook component (sky blue). The hexagonal lattices indicate the positions of the MCP trimers. Individual trimers are labeled P1 to P6 in the asymmetric unit of pentasymmetron, and T1 to T40 in the asymmetric unit of trisymmetron. (C) Three lattice units, Lattice-A, B, and C (red, light blue, and green), which mainly form the mCP network in the trisymmetron under the MCPs, are extracted.

In trisymmetron, the mCP network was formed with a combination of three different lattice components named Lattice lane-A, B, and C (red, light blue, and green in Fig. 2C). These are tandemly connected and repeatedly surrounding the 3-fold axis with the order of A, B, A, and B, from center to sphere (Fig. 2B right). The lane of Lattice-C component enclosed these lattice lines (Fig. 2C). The Lattice-c lines near the corner of trisymmetron were deformed with an interference of the pentasymmetron components (Fig. S2). It shows a novel mCP network forming a large trisymmetron of the giant virus. At the interfaces of pentasymmetron and trisymmetron, lattice lines of Glue component (yellow in Fig. 2) connected each other, and further Hook component existing in the middle of the Glue component’s lattice lines connected the viral capsid to IM (sky blue in Fig. 2)

### Medusavirus MCP structure

The amino acid sequence of medusavirus MCP showed higher homology to that of PBCV-1’s MCP. The MCP was further predicted to have the same “double jelly roll” motif, as a result of a pairwise alignment with PBCV-1’s MCP (Fig. S3). The molecular model of the MCP was generated by AlphaFold2 (14), and the resulting model was fitted into the cryo-EM map and modified based on the MCP density using COOT (15, 16) and Phenix (17). As a result, the MCP model of medusavirus fitted the cryo-EM map well, exhibiting that the outer loops 1 composed of DE-1, part of DE-2, and HI-1 on the first jelly roll motif (Fig. 3A and B). Additionally, the outer loops 2 consisted of the remaining portions of DE-2 and HI-2 on the second jelly roll motif. These loops form large density clusters on the MCP trimer. As described below, these clusters of the outer loops (Fig. 3A and B) and the unique third helix in FG-1 loop (Fig. S3) showed different interactions with different types of spikes (Fig. 4). Further, the MCP labeled P4 was confirmed to be rotated by 60° relative to other MCPs contained in the asymmetric unit of pentasymmetron (Fig. 3C), suggesting to keep a spiral assembly pathway of the capsid as reported in other giant viruses (18).

**Figure 3.**
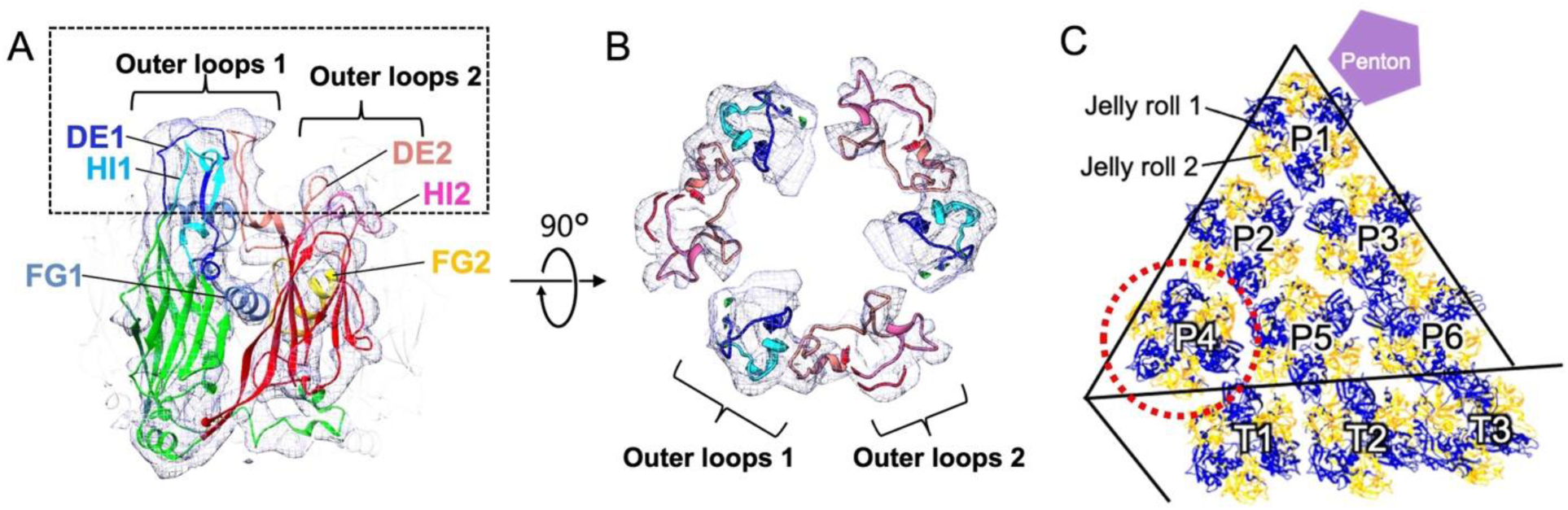
Structure of medusavirus MCP. (A) Cryo-EM map of the MCP monomer and the predicted model fitted. The outer loops of the model are colored as follows: DE1 (blue), FG1 (sky blue), HI1 (light blue), DE2 (salmon pink), FG2 (yellow), and HI2 (pink). DE1, HI1, and a part of DE2 form the outer loops 1 on the first jelly roll motif, and the remaining portions of DE2 and HI2 on the second jelly roll motif form the outer loops 2. (B) Top view of the MCP trimer at dashed box area in A. (C) MCP labeled P4 (dotted red circle) is rotated by 60° relative to other MCPs contained in the asymmetric unit of pentasymmetron.

**Figure 4.**
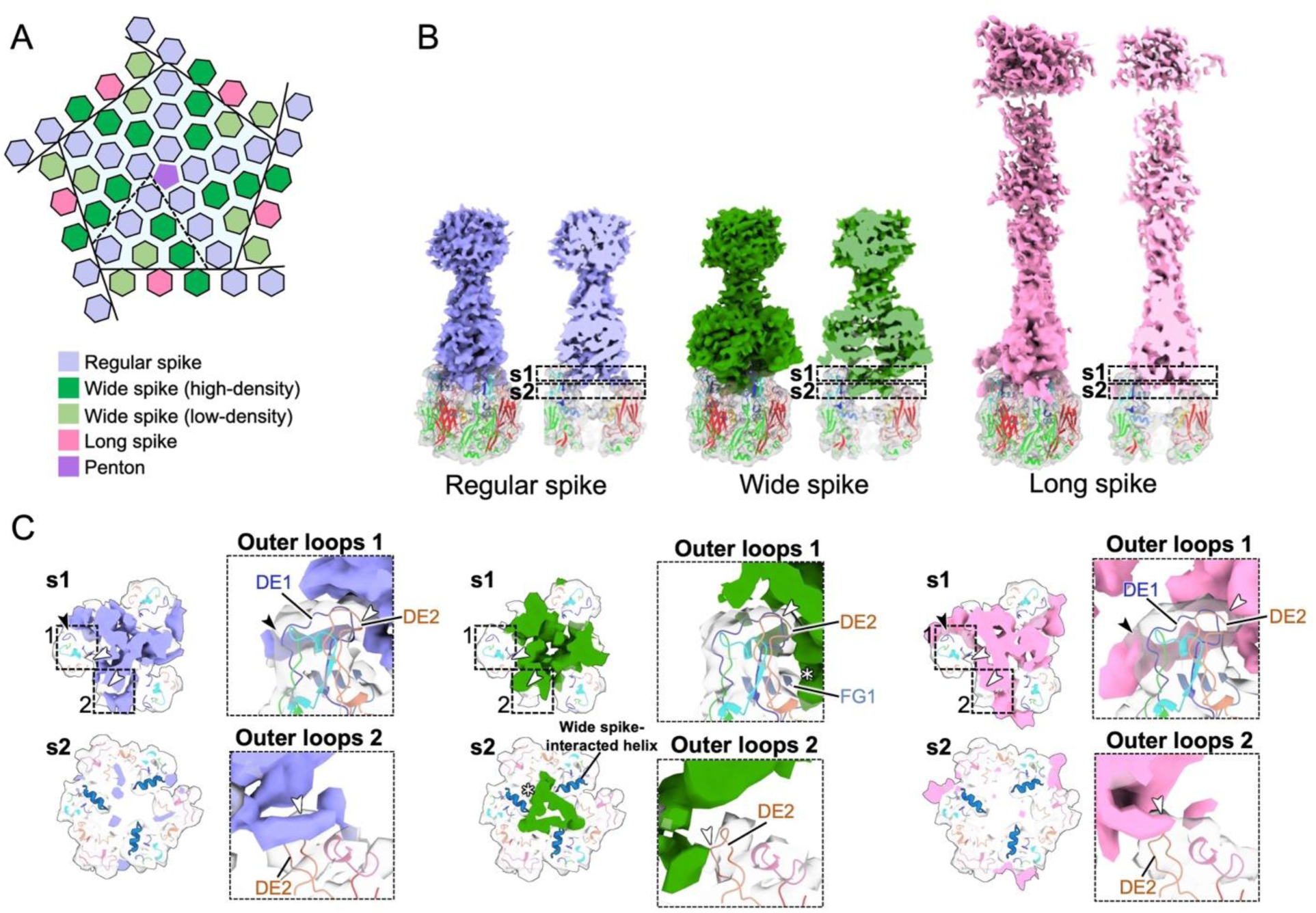
Three types of spikes around 5-fold axis and their interactions with MCPs. (A) Schematic diagram of capsomers (hexagons) arranged around the penton (pentagon) at the 5-fold axis. Each capsomer is colored differently depending on the spike types (B). (B) Structures of the three types of the spikes on capsomers (white). Regular spike (light purple), wide spike (green), and long spike (pink) are indicated. (C) Sliced view of each type of spike, cutoff at dashed boxes a and b in panel B. Focused views of the dashed boxes 1 and 2, indicating interactions between the outer loops 1 and 2 of each MCP and each spike.

### Three types of spikes

As we reported previously (4), medusavirus capsid was decorated with three types of spikes, named regular, long, and wide spikes (Fig. 1D), and we now found that these spikes interact differently with MCPs (Fig. 4). The regular and long spikes were similarly interactions with MCPs, where the root of the spikes interacted with MCP through DE1 and a part of DE2 in the loops 1 and the remaining portion of DE2 in the loops 2 as surrounding these two outer loop clusters (black and white arrows in Fig. 4C right and left panels). Resultantly, interactions were occurring over a wider range of areas. In contrast, the wide spikes interacted with MCPs as sitting on them and interations were formed through a part of DE2 (black arrows) and FG1 (white asterisk) in the loops 1 and a limited part of DE2 in the loops 2 (Fig. 4C middle panel). Interestingly, there were three helices in FG1 loop in MCP of medusavirus, which are only two in those of PBCV-1, ASFV, and SGIV. The third helix uniquely interacted with the wide spikes (white asterisk in Fig. 4C middle panel) (“wide spike-interacted helix” in Fig. S3), but the functions in the different types of spikes are not clear at present.

### The minor capsid proteins around the 5-fold axis

At the 5-fold vertices, the composition of the mCPs varied dramatically depending on the particle type and the presence or absence of internal membrane (IM). In the DNA-Full particle, PC-IV was lost compared to the DNA-Empty particle, even if IM is present. This may be caused by a shrinkage of the IM due to DNA packaging (Fig. 5A and B). In DNA-Empty particle, there were two types of vertices in the capsid. The first has an IM under the vertex, and the second has no IM due to the open structure of the IM (Fig. 5B and C). In addition to the PC-IV, PC-III was also lost on the open structure of IM in DNA-Empty particles. However, the PC-III was recovered in the DNA-Full particle by closing the open structure of the IM.

**Figure 5.**
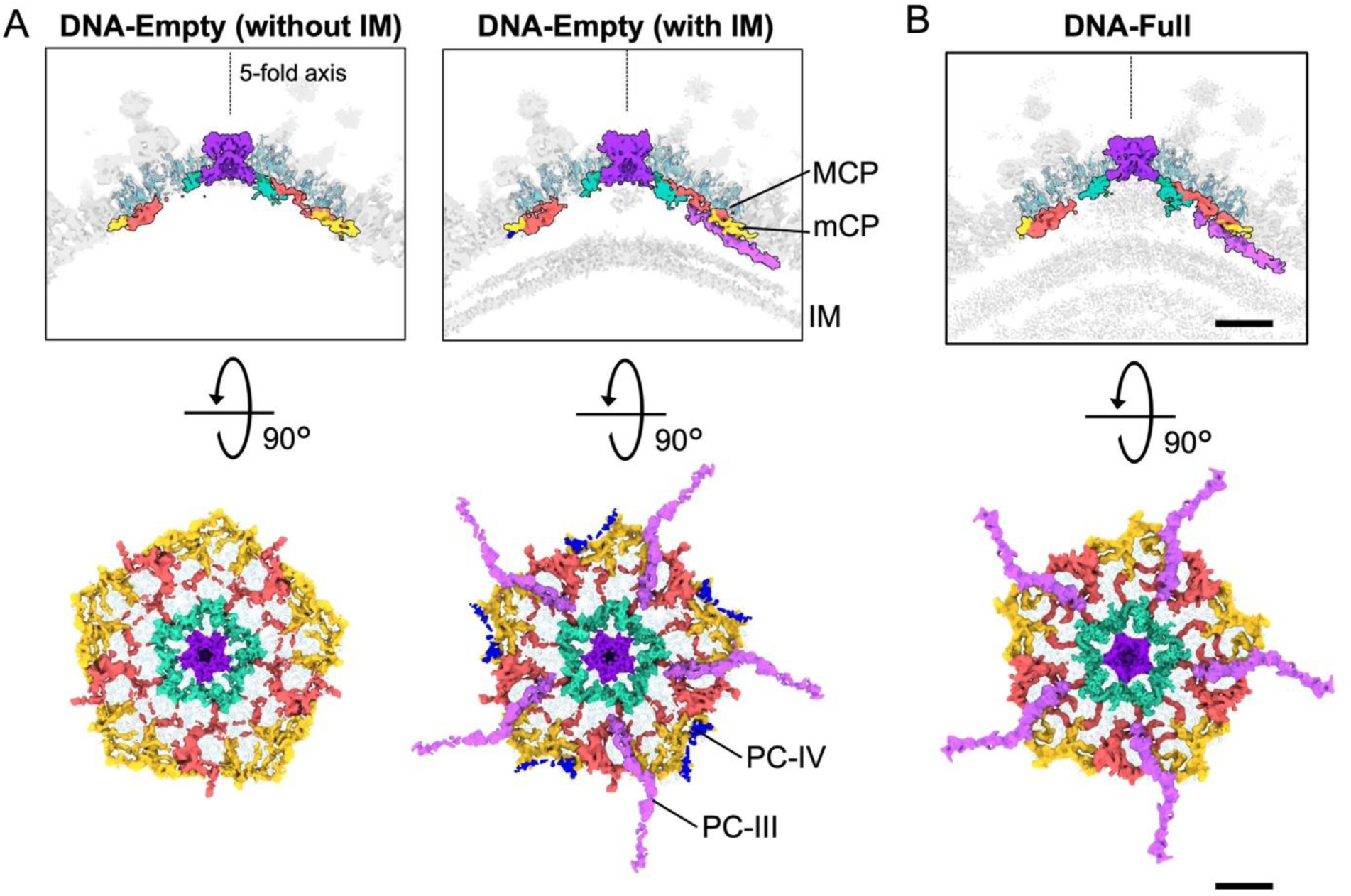
The minor capsid proteins in the DNA-Empty without IM, the DNA-Empty with IM, and the DNA-Full particles. (A and B) Vertical slices of the 5-fold maps colored corresponding to Fig. 2 (top panel), and the minor capsid proteins viewed from inside the capsid (bottom panel). The maps show the DNA-Empty’s blocks with and without IM (A), and the DNA-Full’s block (B). Scale bar = 100 Å.

### Penton protein in DNA-Empty and DNA-Full particles

The structural changes depending on the particle types were also identified at the penton on the 5-fold vertices (Fig. 6). The density of the penton in DNA-Full was shorten 14 Å compared to the DNA-Empty, indicating that the lower part of the penton protein (pink in Fig. 6B) was lost. This may be caused by a shrinkage of the IM due to DNA packaging.

**Figure 6.**
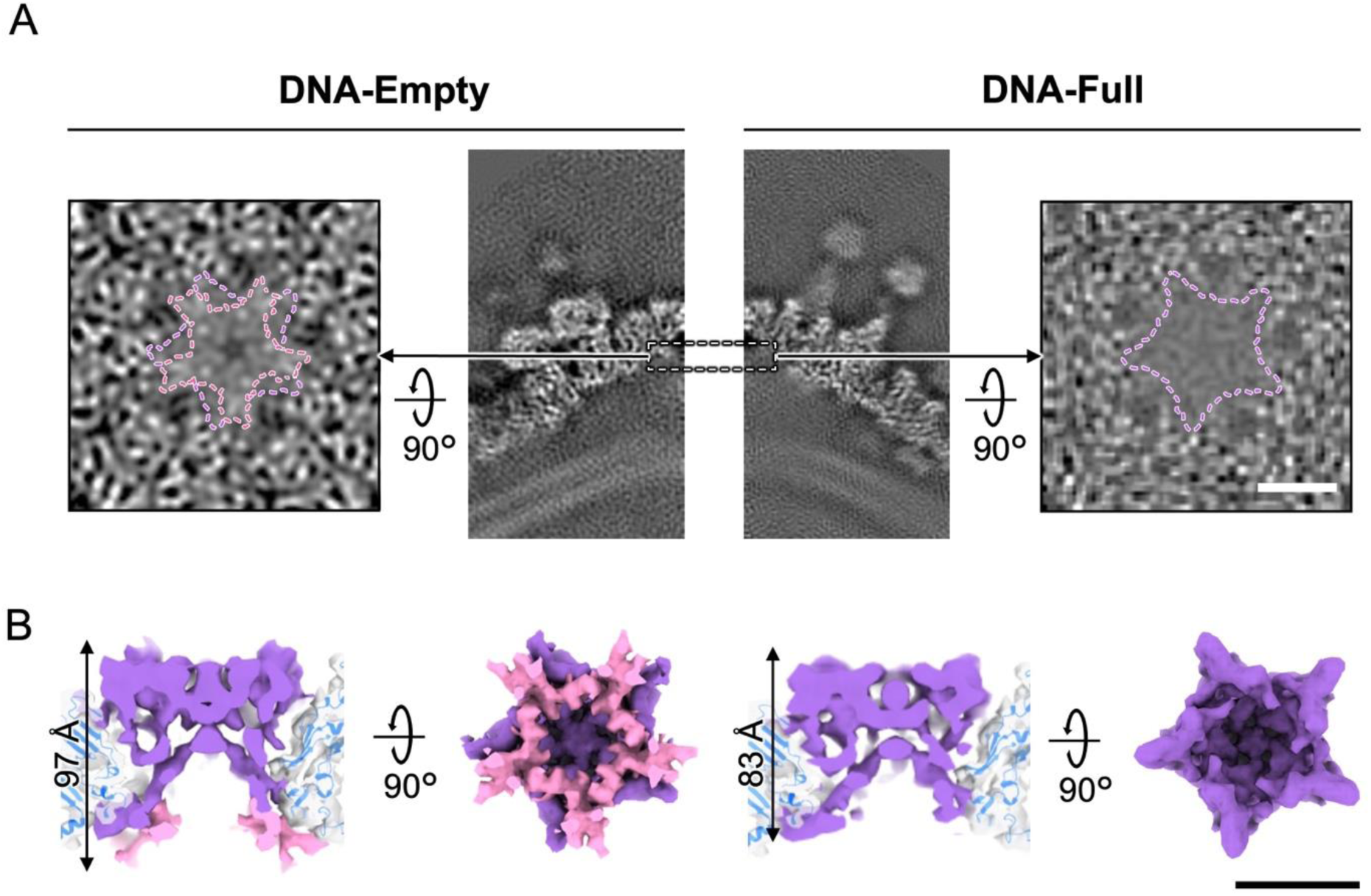
Structural changes of the penton protein between DNA-Empty and DNA-Full particles. (A) Horizontal and vertical slices around the penton in the DNA-Empty (left panels) and DNA-Full (right panels) particles viewed from the inside of the capsid. (C and D) Structures of the penton in the DNA-Empty (C) and the DNA-Full particles (D). The differences are colored by pink. Scale bars = 50 Å.

### Hook components along 2-fold axis

The mCP components named “hook” were located symmetrically along the trisymmetron interface, that bridge the capsid and the IM (yellow arrows in Fig7A to C). The hook components bound in the middle of three MCPs labeled T8, T9, and T20 through the Glue and Lattice-C mCP components (Fig. 7D). These hook components function to maintain the IM structure in both DNA-Empty and DNA-Full particles (Fig. 7B).

**Figure 7.**
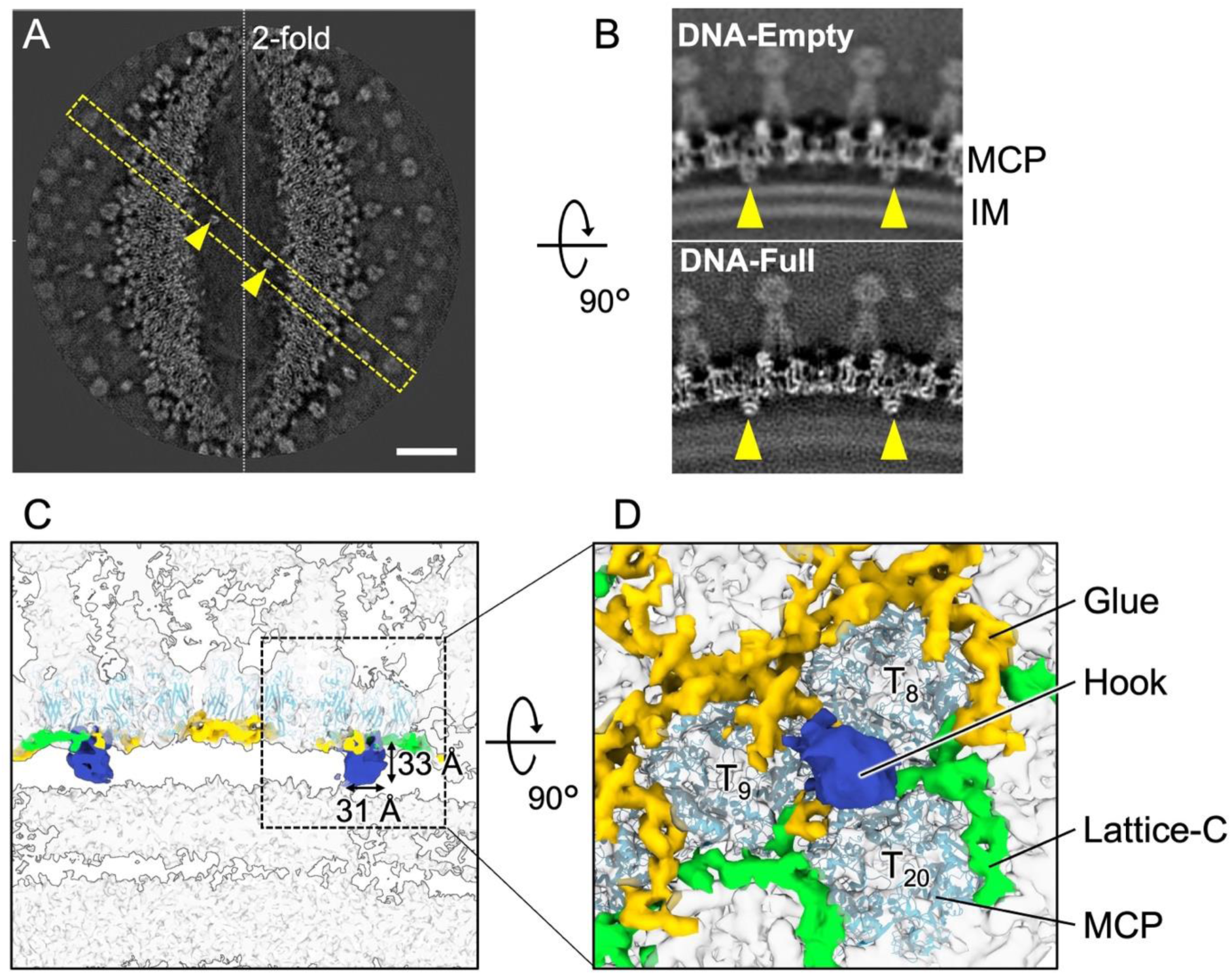
Hook structures connecting capsid and IM along the 2-fold axis. (A) A horizontal slice of the 2-fold cryo-EM map cutoff between capsid and IM. Scale bar = 200 Å. (B) Vertical slices of the DNA-Empty and -Full maps at the yellow dashed box in A. Hook structures are indicated with arrowheads. (C) A vertical slice of the cryo-EM map of the Hook structures along the 2-fold axis, which are colored blue. (D) Focused horizontal view of the dashed box in C, showing the location of the Hook structure in the mCP network.

## Discussion

For medusavirus capsid, the local resolution calculations showed that the regions of the outer spikes were worse than those of other MCPs and mCPs, probably due to flexibility (Fig. S1). The intensities of these spikes were various even for the same type of spikes. For example, the wide spikes on three MCP trimers labeled P3, P6 and T3 showed strong intensity, whereas the wide spikes on two MCP trimers labeled P5 and T1 showed weak intensity (Fig. S4A). In contrast, all long spikes showed strong intensity. These strong intensities in long spikes were thought to be caused by interactions with the neighboring spikes, even though they were expected to be more flexible compared to the shorter wide and regular spikes. There, the wide spikes on the MCP trimers labeled P3, P6 and T3 showed interactions between them at several points (black and white stars in Fig. S4A and C). The long spike on the MCP trimer labeled T2 also showed an interaction with the regular spike on the MCP trimer labeled T14 (black asterisk in Fig. S4A and C). These molecular interactions contribute to the stability of these spikes and cause a strong intensity in the cryo-EM map.

On the other hand, no such interaction was observed between the wide spikes on the MCP trimers labeled P5 and T1 (Fig. S4A and C). To compare details, we defined a central axis on each MCP trimer attached to the wide spike, and measured the tilt angles between each (Fig. S4B). Under the dense wide spikes, the axes of each MCP trimer were not parallel but tilted inward at angles of 13.2° between P3 and P6, and 3.0° between P6 and T3. In contrast, under the weak wide spikes, the axis of each MCP trimer was tilted outward at angle of 6.2° between P5 and T1, while the axis of each MCP trimer between the dense wide spike labeled P3 and the weak wide spike labeled P5 was tilted inward at angle of 17.9° (Fig. S4B). Fitting a typical cryo-EM map of wide spike to each location of the weak wide spikes on the MCP trimers labeled P5 and T1 resulted in an overlap (red area in Fig. S4B). This indicated that the angle between the MCP trimers labeled P5 and T1 was not appropriate to form a molecular interaction between wide spikes. Therefore, the weak density of these wide spikes may indicate partial occupancy of these spikes or the result of a distortion of the spikes.

We confirmed that the MCP trimer located at the corner of the asymmetric unit in pentasymmetron is rotated by 60° (Fig. 3C) similar to other giant viruses (6). Further, we investigated the tilt angle of each MCP trimer in the asymmetric unit of the pentasymmetron compared to the MCP trimer labeled P1 (Fig. S5). The results showed that the tile angles of each MCP trimer are quite similar to those of other giant viruses that don’t have spikes, strongly suggesting that MCP trimers are first assembled to build the capsid, and spikes are then decorated on the MCP array. Wide and long spikes appeared to decorate the higher tilted MCP trimers compared to those of regular spikes (Fig. S5), but some of these spikes are finally stabilized by interactions with neighboring spikes. In the case of wide spikes, the unique third helix in the FG1 loop may help to connect the spikes even in the high tilt condition of the MCP trimers (Fig.4 and S3).

Previous studies have reported that the open structures in the internal membrane (IM) are finally closed and the particle size is shrunk ∼1 nm in the morphological change from DNA-Empty to DNA-Full particles (4). In this study, we observed novel structural changes, in which PC-III, IV and penton interacted with IM were deformed or lost between these (Fig. 5 and 6). On the other hand, despite the drastic changes of IM during the maturation process, the distance between the capsid and IM via hook was maintained (Fig. 7B). This suggests that the hook component plays an important role in maintaining the morphology of the IM through the maturation process.

Two membrane proteins can be predicted for medusavirus (encoded by the orf226 and orf329 genes) (1). The membrane protein encoded by the orf226 gene shows homology to that of the *Marseilleviridae*. On the other hand, the myristoylated membrane protein encoded by the orf329 gene shows homology with VP88, the anchor protein of SGIV (8), by PSI-BLAST search (19). The VP88 is an mCP that contacts the inner membrane at the trisymmetron-pentasymmetron interface in SGIV. As comparing to VP88, medusavirus’s myristoylated membrane protein would be expected as a candidate for PC-III, IV or hook. Higher resolution of these densities will be necessary to identified the homologous mCP in the future.

MCP of medusavirus exhibits high homology with other giant viruses such as PBCV-1 and SGIV (Fig. S4A), and the orientation pattern of the MCP trimer on the capsid surface is also similar to that of other giant viruses (Fig. 3C). However, except for the VP88 of SGIV discussed above, homologs of mCPs were not identified in the medusavirus genome. Tape-measure protein is a mCP previously commonly identified in several icosahedral giant viruses of PBCV-1 and ASFV (7 and 13) in addition to bacteriophages of PRD1 and Bam35 (20, 21), and proposed to determine capsid size by connecting 5-fold vertices (18). However, we did not find the genome and the corresponding density in the medusavirus capsid even though the resolution is not enough to individually elucidate this. CIV (22), tokyovirus (12), and melbournevirus (11) were reported to have no tape-measure protein. In these giant viruses including medusavirus, another mechanism may exist to determine the large capsid size.

Concerning the currently limited resolutions of the medusavirus maps, the FSC curves drops down at 7.3-9.9 Å resolution even after the block-based reconstructions (Fig. S1I). The facts suggest that no higher-resolution information is included in the EM images, despite the sufficiently high image magnification. The local resolutions showed that the resolutions are significantly reduced the outer spikes (Fig. S2), suggesting that the flexibility of the outer spikes influences the entire capsid stability. Apply a local mask to the spikes would somewhat improve the resolution of the capsid part, but the instability of the capsid through the maturation process may have a key role for the special medusavirus life cycle (4).

In this study, we revealed some dynamical structural changes of the mCP components in different types of medusavirus particles which show each virus maturation stage (4). There, some mCP components are lost in advancement of maturation, suggesting that these components temporally exit during the maturation process and didapper after the maturation. These temporal mCP components were reported for the first time in giant viruses, as medusavirus is also released as immature particles in addition to mature particles. Finding of these temporary components is important to clarify the particle maturation process of giant viruses. Medusavirus is unique to be able to investigate the immature particles in addition to the mature virion. Further structural investigation of the virus particles will reveal the mechanism to form the large viruses in the future.

## Materials and Methods

### Medusavirus growth and purification

Medusavirus was grown and purified by infecting *A. castellanii* strain Neff as previously described (4). Briefly, amoeba cells were cultured at 26°C in flasks containing 100 mL of peptone-yeast-glucose (PYG) medium, and medusavirus was harvested 3 days post infection. Amoeba cells and cell debris were removed by centrifuging at 800× g for 5 min at 24°C, and the supernatant was centrifuged at 8,000× g for 35 min at 4°C to pellet the viral particles. The viral particles were suspended in phosphate-buffered saline (PBS), and purified using a 0.45-μm filter (Millex-AA; Merck Millipore, Darmstadt, Germany). The filtered viral particles were centrifuged at 8,000× g for 35 min at 4°C, and then resuspended in PBS. This process was repeated multiple times to obtain a sufficient amount of medusavirus particles.

### Cryo-EM Data Collection and Processing

The data collection and image processing procedures are described previously (4). In brief, a 2.5 mL suspension of purified medusavirus particles was applied to a glow-discharged Quantifoil grid (R1.2/1.3 Mo; Quantifoil Micro Tools GmbH, Germany). The grid was blotted with filter paper (blotting time, 7 s; blotting force, 10 in the Vitrobot setting) and plunge-frozen at 4°C under 95% humidity using a Vitrobot Mark IV (Thermo Fisher Scientific, USA). The frozen grid was imaged using a Titan Krios G3 microscope at 300 kV equipped with a Falcon III detector. The datasets were recorded at a nominal magnification of ×22,500, corresponding to 3.03 Å per pixel on the specimen. A low-dose method (exposure at 10 electrons per Å^2^ per second) was used, and the total number of electrons accumulated on the specimen was ∼30 electrons per Å^2^ for a 3 second exposure time. A GIF Quantum energy filter (Gatan, Inc., USA) was used with a slit width of 20 eV to remove inelastically scattered electrons. Individual micrograph movies were subjected to per-frame drift correction by MotionCor2 (23), and the contrast transfer function parameters were estimated by CTFFIND4 (24). Subsequently, 4,625 DNA-Empty and 7,038 DNA-Full particles were manually selected and extracted from 2,084 motion-corrected images using RELION3.0 software (25). 4,551 DNA-Empty particles and 6,981 DNA-Full particles were selected from 2D classification and used for 3D reconstruction with imposed icosahedral symmetry. The resolutions of the entire capsids of the DNA-Empty and -Full particles were estimated at 21.5 Å and 19.5 Å using the 0.143 gold standard FSC criterion (26). The reconstructed 3D maps were further improved using a block-based reconstruction strategy (10) to account for defocus gradient and particle flexibility, and resulted in final resolutions of 7.3-9.9 Å. The cryo-EM map of the major capsid proteins (MCPs) was interpreted by fitting the molecular model generated by AlphaFold2 (14). The fitted model was manually fixed by COOT (15) and refined by PHENIX (17).

## Data availability

The cryo-EM maps have been deposited in the EMDB with the accession codes EMD-39292 (5-fold block of the DNA-Full particle), 39293 (5-fold block of the DNA-Empty particle with internal membrane), 39294 (5-fold block of the DNA-Empty particle without internal membrane), 39295 (3-fold block of the DNA-Full particle), 39296 (3-fold block of the DNA-Empty particle), 39297 (2-fold block of the DNA-Full particle), 39298 (2-fold block of the DNA-Empty particle), respectively.

## Acknowledgments

We thank for Dr. Raymond Burton-Smith for assistance with block based reconstruction claculations. This study was supported by MEXT/KAKENHI under grants JP17H05825 and JP19H04845 to K.M., JSPS/KAKENHI under grant 20H03078 to M.T., BINDS from AMED under grant JP18am0101072 (support number 1162) to K.M., Joint Research of ExCELLS under grant 20-004 to K.M., and the Cooperative Study Program of the National Institute for Physiological Sciences under grant 20-239 to M.T.

## Author contributions

K.M. conceived the project. M.T. provided the medusavirus sample. C.S. and R.W. prepared the cryo-EM sample and collected the cryo-EM data. C.S., R.W., and K.M. processed the cryo-EM data. R.W. prepared the figures. R.W. and K.M. wrote the main manuscript text. All authors reviewed the manuscript.

