## Supplemental figures for "Subnanometer structure of medusavirus capsid during maturation using cryo-electron microscopy"

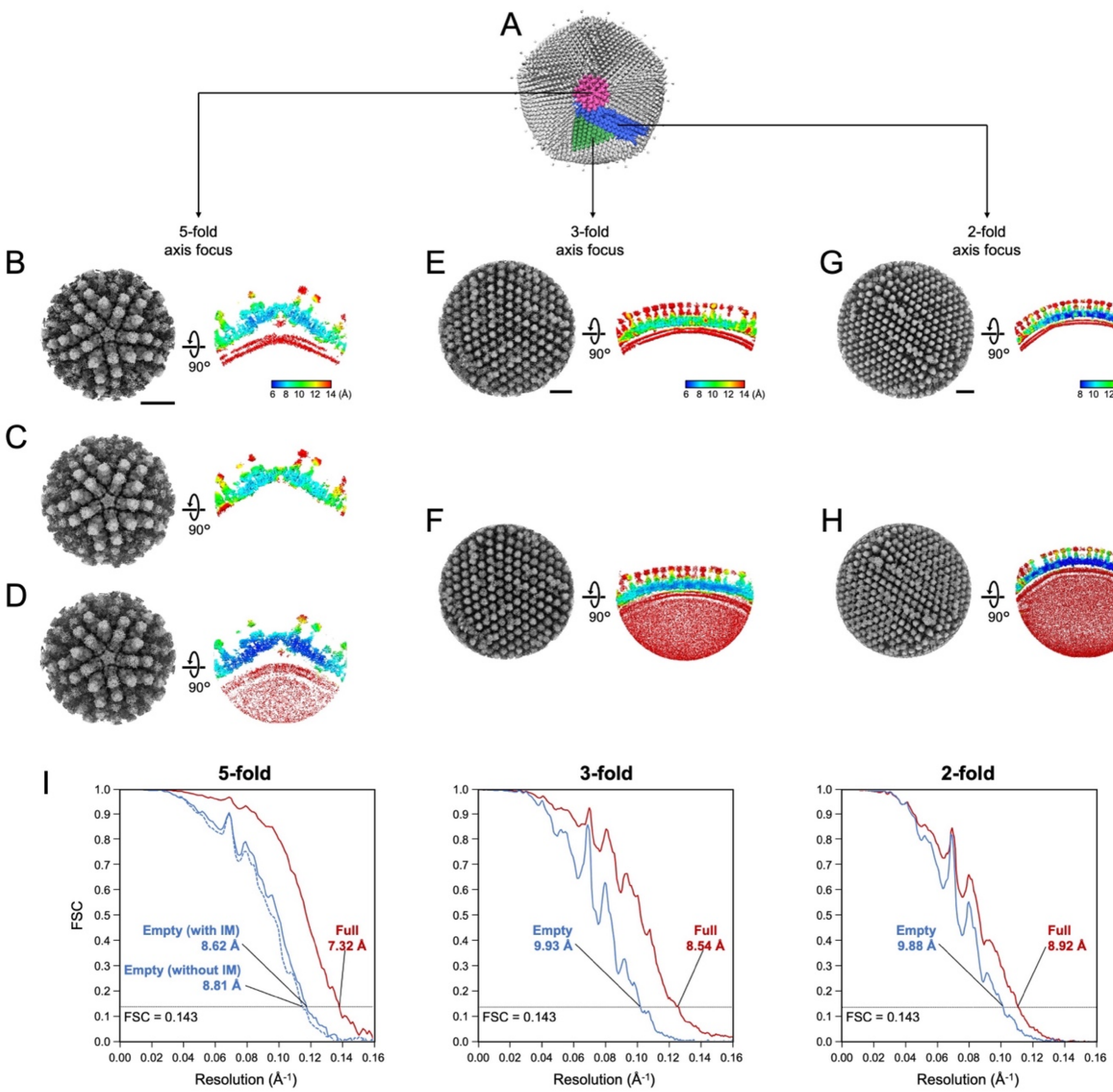


**Fig. S1 Single particle image processing flowchart for the block-based reconstructions**

(A) Blocks centered on the 5, 3, and 2-fold axes in the medusavirus capsid are colored pink, green, and blue, respectively. (B, C and D) Cryo-EM maps of the 5-fold block (left panel), and the vertical slices of the maps along the 5-fold axes colored by local resolution (right panel). The maps show the DNA-Empty capsid with IM (B), the DNA-Empty capsid without IM (C), and the DNA-Full capsid (D). (E and F) Cryo-EM maps of the 3-fold block (left panel), and the vertical slices of the maps along the 3-fold axes colored by local resolution (right panel). The maps show the DNA-Empty capsid (E), and the DNA-Full capsid (F). (G and H) Cryo-EM maps of the 2-fold block (left panel), and the vertical slices of the maps along the 2-fold axes colored by local resolution (right panel). The maps show DNA-Empty capsid (G), and DNA-Full capsid (H). Scale bars = 200 Å (I) Gold standard FSC curves of 5-fold, 3-fold, and 2-fold, respectively. DNA-Full and DNA-Empty with and without IM are included by different colors.

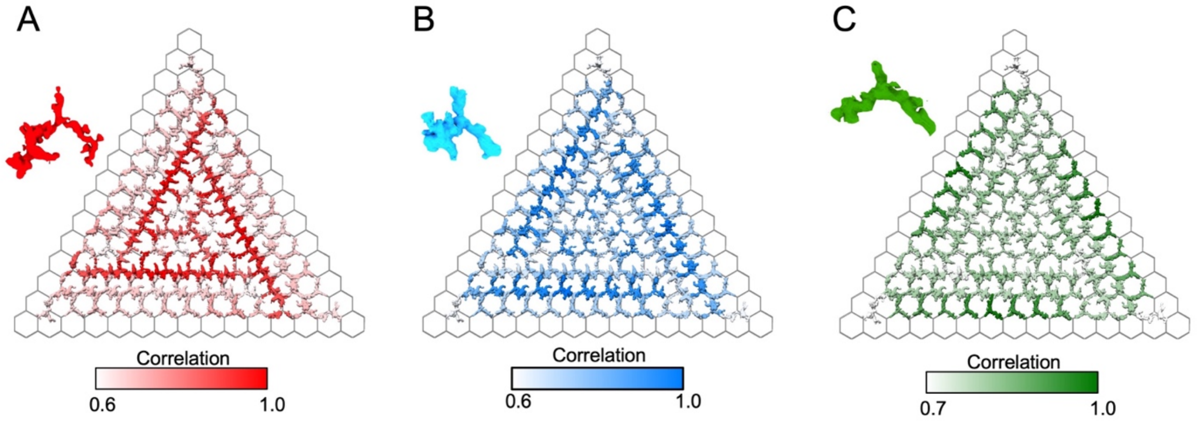


**Fig. S2 Three lattice units and their correlation in the mCP network**

The correlations of three lattice units (Lattice-A, B, and C) (red, light blue, and green) with the mCP networks under trisymmetron were calculated, and the correlation values are colored.

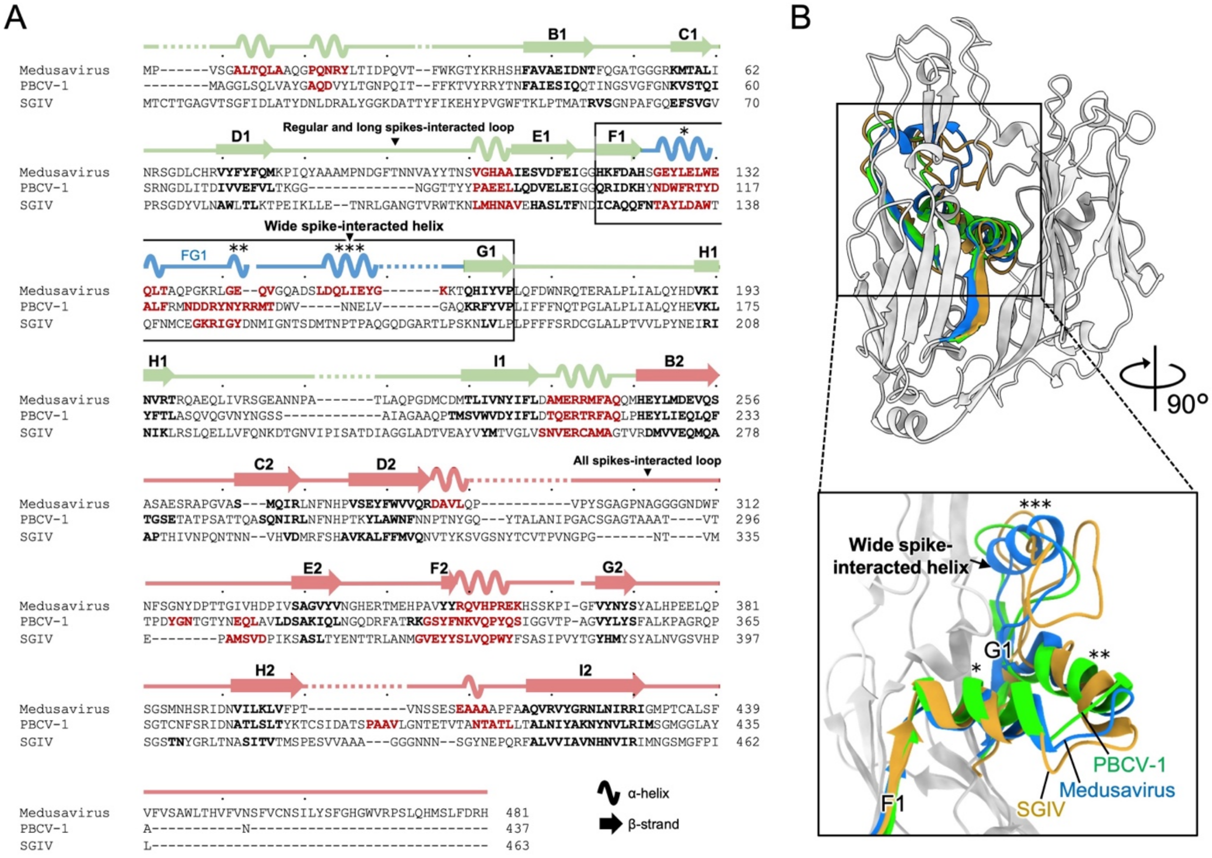


**Fig. S3 Sequence alignment of MCP and comparison of the helix in FG1 loop**

(A) Multiple alignment of MCP sequences among MedV, PBCV-1, and SGIV.

(B) Superposition of MCP ribbon diagrams of medusavirus (sky blue), PBCV-1 (light green), and SGIV (orange). Focused views of the dashed box for comparison of FG1 loops among the viruses.

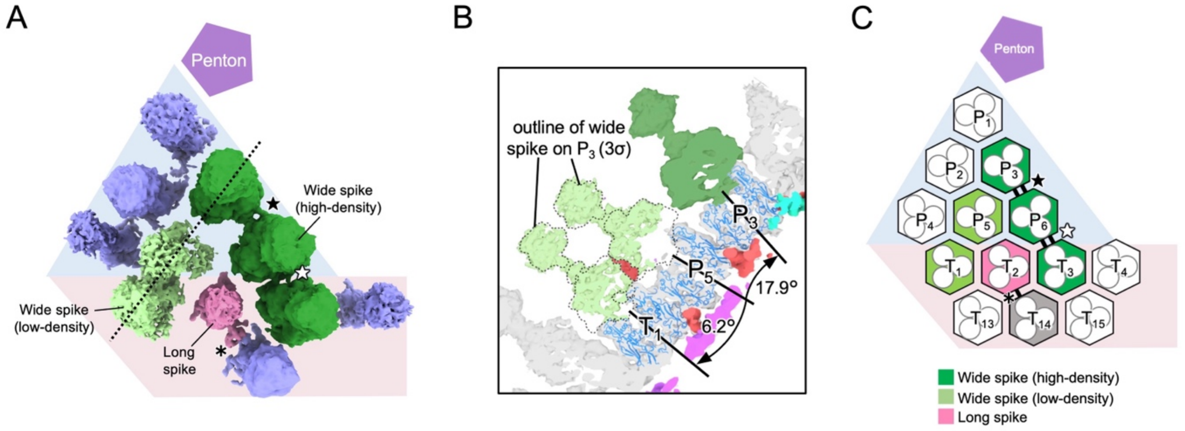


**Fig. S4 Cryo-EM maps of each spike, and their interactions**

(A) Cryo-EM map of each spike around the asymmetric unit of pentasymmetron. Each spike is colored corresponding to Fig. 4A. Interactions between spikes are marked with black stars, white stars, and asterisks. Contour level = 1σ

(B) Vertical slice of the cryo-EM map at the dashed line B in A. The radiation angles of the spikes are indicated.

(C) Schematic diagram showing three spike types and interactions between them.

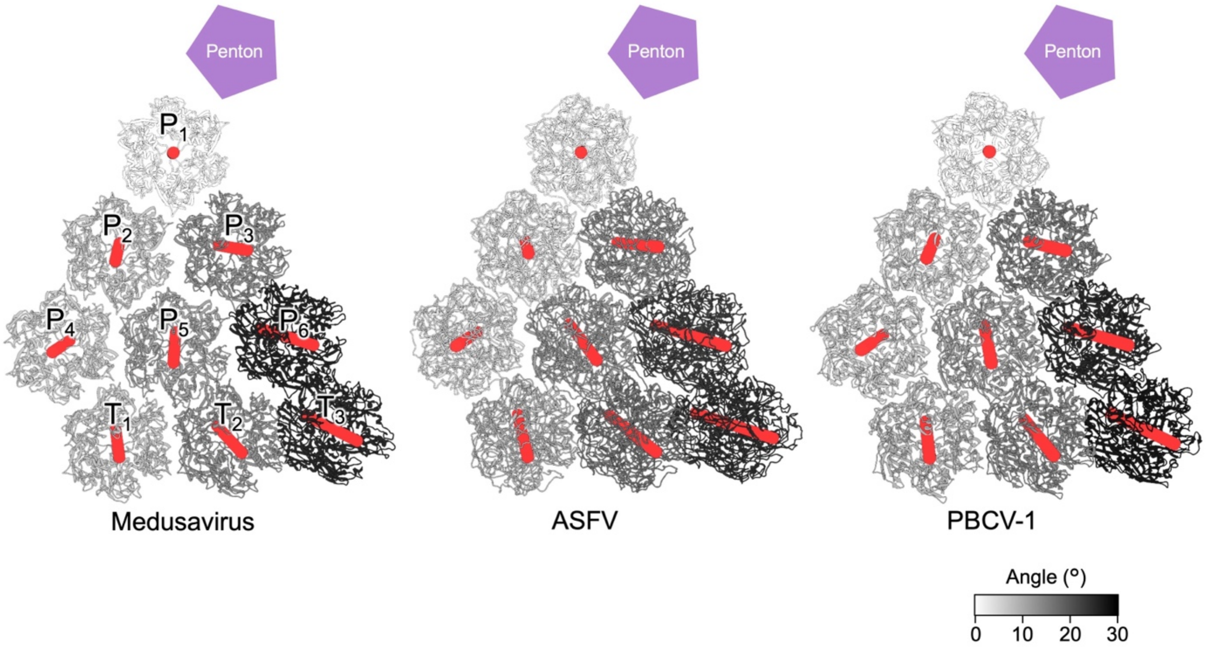


**Fig. S5 Relative tilted angles of each MCP compared with the nearest MCP (P1) adjacent to the penton in different viruses**

The MCP capsomers (P1 to P6, and T1 toT3) are colored according to the tilted angles indicated.
